# Universal Study Design for Instrument Changes in Pharmaceutical Release Analytics

**DOI:** 10.1101/2024.12.11.627881

**Authors:** Anne B. Ries, Maximilian N. Merkel, Kristina Coßmann, Marina Paul, Robin Grunwald, Daniel Klemmer, Franziska Hübner, Sabine Eggensperger, Frederik T. Weiß

## Abstract

Instrument changes in analytical methods of pharmaceutical quality control are required to maintain release analytics over decades, yet typically pose a challenge.

We designed an efficient instrument comparability study to gain a comprehensive understanding of potential performance differences between instruments and therefore rationalize the risk assessment and decision process for a path forward. The results may either point out whether a full or partial re-validation is necessary or whether a science-based bridging can be pursued based on the data generated in the study. The study design is universally applicable to a substantial range of release analytical methods. In a straightforward setup of two experiments with the new instrument, a statistically meaningful data set is generated for comparison with available historical or validation data of the original instrument.

In a Good Manufacturing Practice (GMP) environment, we realized the study design a first benchmark in imaged capillary isoelectric focusing (icIEF) analytics, comparing the ICE3 and Maurice C instruments. The core-study confirmed equal or better performance of Maurice C in all parameters and serves as a basis for seamless continuation of release measurements on Maurice C.

## 1 Introduction

Analytical methods in pharmaceutical quality control (QC) need to be maintained for decades to guarantee release analytics and continued or ongoing process verification over the life cycle of commercial products [1–4].

Upon e.g., an instrument update, product specific analytical procedures validated on the original equipment must be moved to the new equipment [2]. The implementation of such technical changes in a regulated environment typically poses a challenge and is of high interest in the respective field [5–7].

Depending on the impact of the technical change on the analytical result, a science-based instrument bridging effort, a partial or a full method re-validation may be considered [2,8–11]. The impact furthermore determines the type of change and regulatory paths to follow when implementing the new instrument [12].

In line with ICH Q2(R2), Q9(R1) and Q14 [9–11], we developed a universal study design to assess instrument comparability comprehensively yet efficiently, to minimize subjectivity, and to provide a science-based decision in the path-forward of instrument modernization in a Good Manufacturing Practice (GMP) environment.

If instrument comparability can be fully confirmed, the study design may well serve as scientific justification to seamlessly continue release analytics on the new instrument.

The proposed study design aims at methods with a graph-based readout, such as capillary-electrophoretic or chromatographic methods, and the underlying concept can be adapted to other methods. Following this design, two experiments on the new instrument are sufficient to generate statistically meaningful data. For evaluation of comparability with the original instrument, no further experiments are required to be carried out, instead, validation and historical data are used.

If multiple products are affected by the instrument change, the product with the highest complexity in readout and sample preparation is used in the comparability study. Thereby, the study results may be considered indicative for further affected products / procedures.

As a benchmark, we applied the study design in a comparability assessment of two imaged capillary isoelectric focusing (icIEF) instruments ICE3 (protein simple) and Maurice C (protein simple) in a GMP environment.

## 2 Theory

In an ideal situation, two technically differing instruments of the same analytical purpose produce identical results. Thus, comparison of the instrument readout is key to understanding potential discrepancies.

To systematically assess comparability of graph-based readouts, such as electropherograms or chromatograms, we considered discrepancies along the x-axis, y-axis, and within the curve integral area (Figure 1). Furthermore, we took the method variance between measurements into account. These considerations led to the identification of seven parameters, namely

- y-axis

1. limit of signal quantitation
2. proportionality between product concentration and observed signal
3. baseline comparability
- x-axis

4. peak position shifts
5. resolution
- integral

6. peak area changes
- measurement variance

7. across different instruments, analysts, days, capillaries/columns.

**Figure 1:**
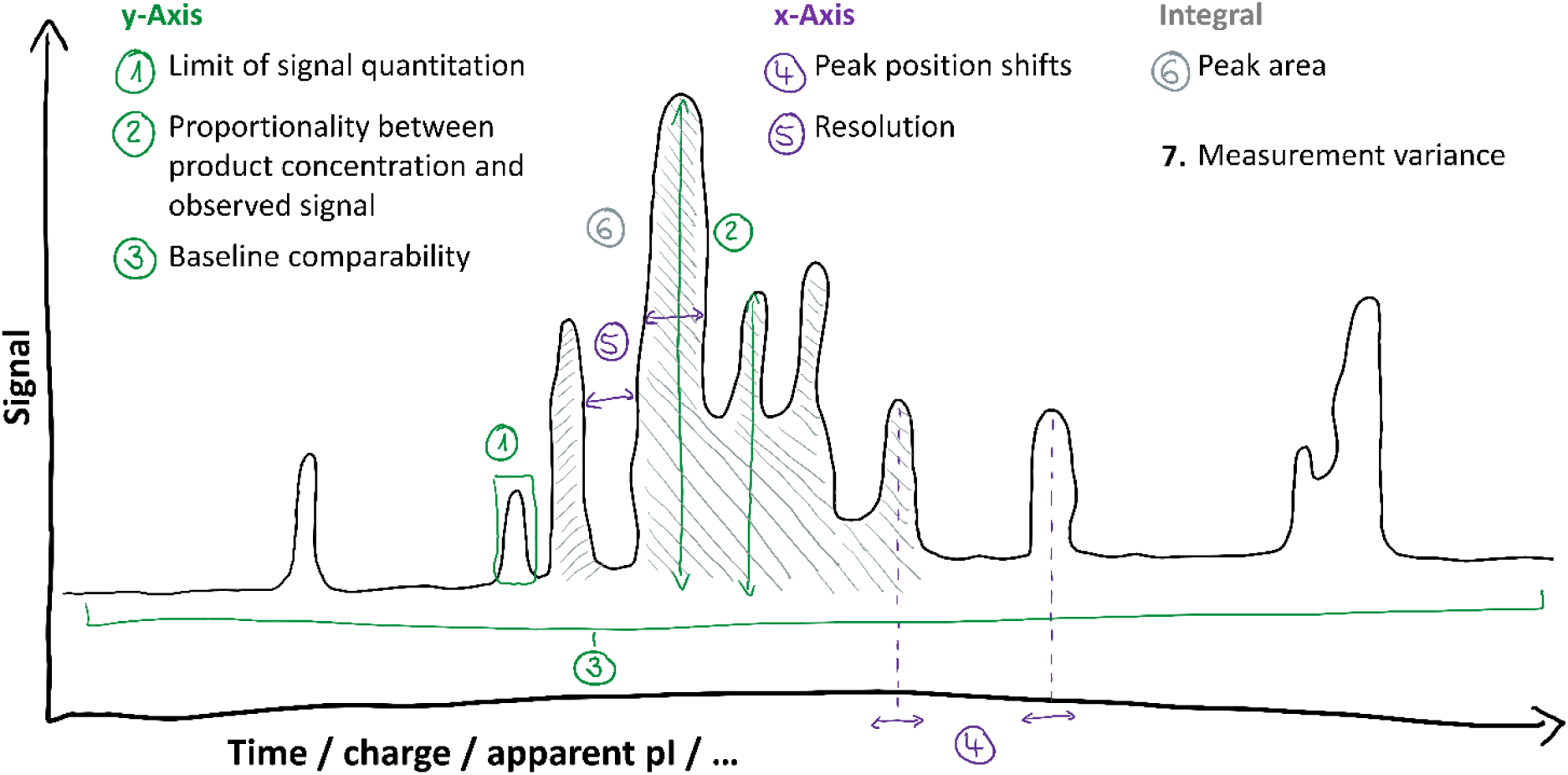
Visualization of graph parameters to be assessed in y-dimension (green labels 1-3), x-dimension (purple labels 4,5) and surface area of the curve integral (grey label 6).

The acceptance criterion for instrument comparability should be the observation of equal or better performance of the new instrument [2]. To set individual acceptance criteria for each parameter, acceptance criteria of the method validation on the original instrument or historical data from the original instrument (such as control chart data) may be used. Visual comparison may be advisable for some parameters.

In the following sections each parameter and setting of respective acceptance criteria are described in detail.

### 2.1 Limit of signal quantitation

To evaluate the comparability of detection sensitivity and signal intensities between two instruments, the limit of quantitation (LOQ) is determined on the new instrument, following the procedure from method validation on the original instrument.

The LOQ value is typically determined as the minimal area that can be quantified. An area is not a pure y-axis parameter and especially in case the LOQ was determined from a peak that is not baseline separated from a neighboring peak, resolution (peak width, peak separation) may impact the LOQ value. To exclude the impact of resolution, instrument comparability can instead be evaluated by assessing LOQ peak signal-to-noise (S/N) values (pure y-axis parameter) and peak area RSD (peak robustness). Both measures are typical acceptance criteria for the LOQ during method validation and are thus likely to be available from the validation on the original instrument.

#### 2.1.1 Statistical evaluation

In case the LOQ value is not impacted by resolution, comparison of LOQ point values may be sufficient. The LOQ on the new instrument should be equal or smaller than on the original instrument.

In case the LOQ value is impacted by resolution, non-inferiority of the LOQ-peak S/N values on the new instrument may be assessed with a one-sided t-test (α=0.05).

The LOQ-peak RSD point value calculated from measurements on the new instrument should be identical or lower than the RSD point value determined during validation on the original instrument.

### 2.2 Proportionality between product concentration and observed signal

To evaluate if sample concentration and detected signal intensity show a comparable proportionality on the new instrument, a response experiment needs to be performed, following the procedure and acceptance criteria of the original method validation on the original instrument.

#### 2.2.1 Statistical evaluation

The acceptance criteria for response in the original validation should be met by the response analysis on the new instrument. Typical criteria may include a limit in correlation coefficient r over the linearity range (e.g. r ≥ 0.98) or in the coefficient of determination R^2^.

In addition to the validation criteria, the validation results may be compared to the results acquired on the new instrument.

### 2.3 Baseline comparability

Graph baselines need to be compared between both instruments, to investigate potential trends in the y-dimension of the graph. The parameter can be evaluated through comparison of blank measurements. Multiple (e.g. three) representative blank measurements from historical data of the original instrument and data acquired on the new instrument may be compared in a direct overlay.

#### 2.3.1 Evaluation

Visual comparison allows identification of potential irregularities or discrepancies in putative method specific blank patterns. Irregularities in the blank measurements of the new instrument should be similar or less pronounced than on the original instrument.

### 2.4 Peak position shifts in x

To evaluate consistency of the peak positions in the x-dimension of the graph, exact x-values of multiple (e.g. three, distributed over the separation range of the product) specific peaks should be determined. The peak position data should consist of at least 12 independent data points collected from a matrix setup (chapter 3.1; Kojima design [13], Table S1).

#### 2.4.1 Statistical evaluation

For the peak positions on both instruments to be consistent, the acquired data from the new instrument should be in a range of ±3 standard deviations (SD) of the historical data acquired on the original instrument (e.g. control chart data). The comparability of both instruments should be statistically evaluated with a two one-sided t-test (TOST), using a significance threshold of α=0.05.

### 2.5 Resolution

To assess graph resolution, peak widths and peak separation on both instruments need to be compared.

Therefore, multiple (e.g. three) representative sample measurements from each historical data of the original instrument and data acquired on the new instrument may be visually compared in a direct overlay.

In case the selected product / analytical procedure provides baseline-separated data, resolution should be determined numerically at multiple (e.g. three) specific positions in the graph. The resolution data should consist of at least 12 independent data points collected from a matrix setup (chapter 3.1; Kojima design [13], Table S1).

#### 2.5.1 (Statistical) evaluation

Visual comparison allows identification of potential discrepancies in peak width and peak separation between both instruments. Resolution on the new instrument should be similar or better than on the original instrument.

In numerical evaluation, for the peak width / separation on both instruments to be consistent, the acquired resolution data from the new instrument should be in a range of ±3 SD of the historical resolution data acquired on the original instrument (e.g. control chart data). The comparability of both instruments should be statistically evaluated with a TOST, using a significance threshold of α=0.05.

### 2.6 Peak area changes

To evaluate consistency of the (relative/total) peak areas (i.e. the curve integral) between both instruments, (relative/total) peak areas of specific peaks or peak groups in the electropherogram should be compared. The peak area data should consist of at least 12 independent data points collected from a matrix setup (chapter 3.1; Kojima design [13], Table S1).

#### 2.6.1 Statistical evaluation

For (relative/total) peak areas on both instruments to be consistent, the acquired data from the new instrument should be in a range of ±3 SD of the historical data acquired on the original instrument (e.g. control chart data). The comparability of both instruments should be statistically evaluated with a TOST, using a significance threshold of α=0.05.

### 2.7 Measurement variance

Measurement variance across different instruments, analysts, days, capillaries/columns indicates whether product analysis has a similar level of precision on both instruments. The measurement variance data should consist of at least 12 independent data points collected from a matrix setup (chapter 3.1; Kojima design [13], Table S1).

#### 2.7.1 Statistical evaluation

For measurement variance on both instruments to be consistent, an SD or RSD analysis may be considered:

SD analysis: The acquired data from the new instrument should have a standard deviation equal or below x·SD of the historical data acquired on the original instrument (e.g. control chart data). An empirically recommendable factor x would be 2 and depends on the sample size. Non-inferiority of the standard deviation on the new instrument may be assessed with a one-sided Chi-Square test (α=0.05).

RSD analysis: The RSD point value of the acquired data from the new instrument should be equal or below y·RSD of the historical data acquired on the original instrument (e.g. control chart data). An empirically recommendable factor y would be 1.5, typically used in acceptance criteria setting for the intermediate precision parameter of method validations.

In addition, the intermediate precision results of the original method validation may be compared to the results acquired on the new instrument.

## 3 Materials and methods

### 3.1 Experimental design

The experimental procedure comprises a response and a matrix setup experiment, from which further parameters can be derived (purple boxes in Figure 2).

**Figure 2.**
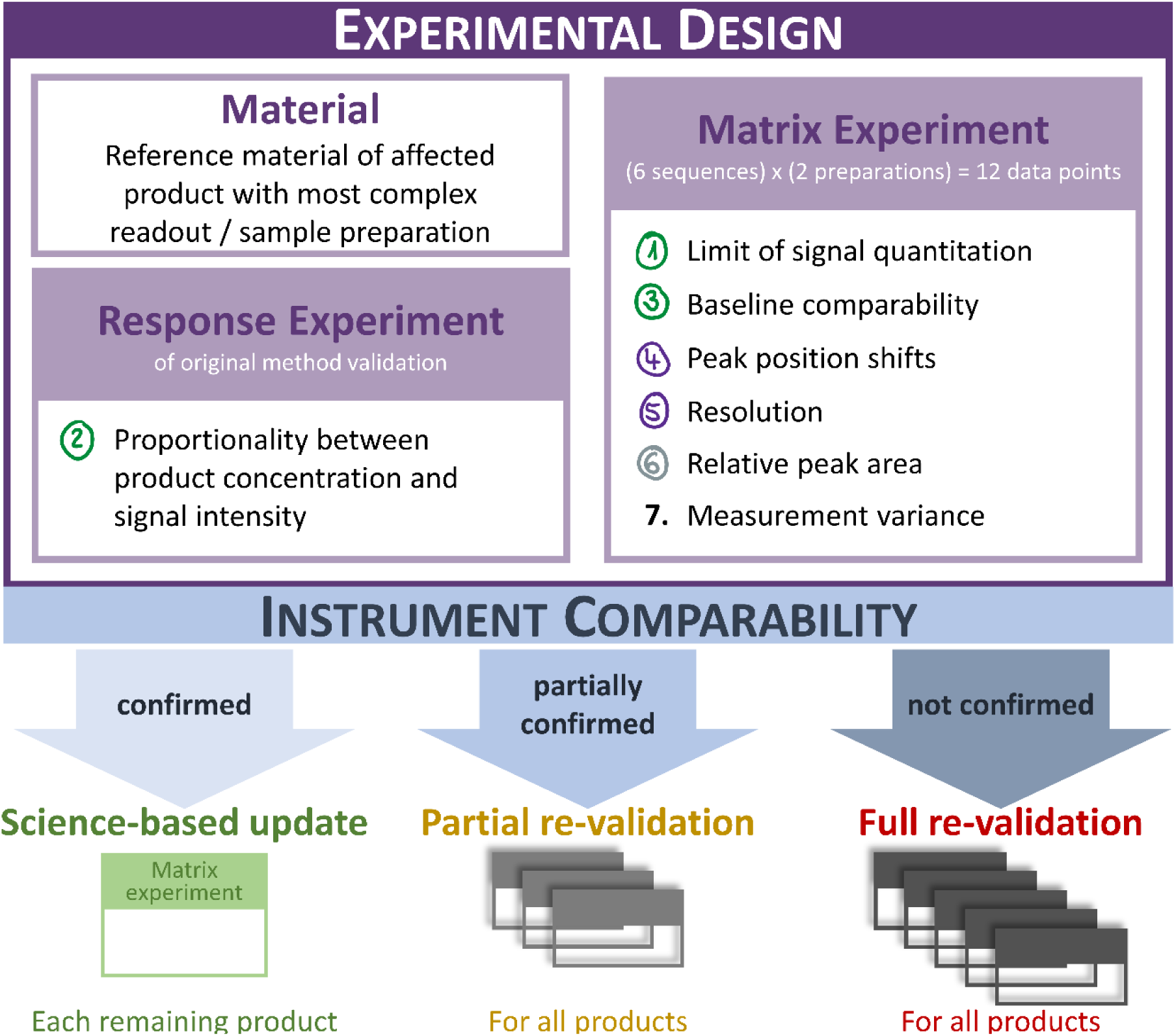
Experimental design of the instrument comparability study: Purple boxes contain experimental setup and derived data of core study, blue box and arrows display potential outcomes. Green, yellow and red headlines depict path forward depending on the identified outcome. Below, the respective analytical effort for each scenario is illustrated.

The sample size of the matrix experiment should at least consist of 12 data points. Other sample sizes may be possible; the DoE set-up for the Kojima design should reflect the strongest variance influencing factors. Furthermore, the sample size should be large enough that given the method variability of the new instrument it is possible to reliably pass the TOST. A power analysis should be done with respect to the new instrument and the acceptance criteria from historical data to get an acceptable passing probability (=power of e.g. at least 90%).

### 3.2 Study material

If multiple products are affected by the instrument change, it is advisable to use the product with the highest complexity in readout and sample preparation in the comparability study. Thereby, the study results may be considered indicative also for other affected products / procedures.

Reference material is the preferable material to carry out comparability experiments on the new instrumentation, as it is advisable to set acceptance criteria for some parameters from historical data collected on the original instrument. In a GMP environment such historical data are typically available from control charts, containing data of reference material over a long period of time. Control chart data provide high statistical power and a good estimate on expected method variation. Furthermore, reference material is typically generated in high amounts and is therefore likely to be available.

In case instrument comparability can be inferred, the study results may well be foundation for the instrument change for all affected products. It is advisable to complement the study with a) a test whether also the product specific SST criteria of the product used in the study are met in the experiments on the new instrument and b) a lean experimental design, investigating the other affected products on the new instrument and investigating whether the product specific SST criteria are met. The matrix experiment is well suited for that purpose (green box in Figure 2).

To ensure system suitability in all experiments, a product-independent and well-studied material is recommended to be used. If available, an instrument specific kit by the manufacturer may be an optimal choice.

In case instrument comparability can only partially or cannot be inferred, the collected data provide insights into the critical discrepancies between both instruments and can serve as a scientific basis for the setup of a partial or full revalidation strategy (Figure 2, yellow and red outcomes).

### 3.3 icIEF instrument comparability study

The study design was applied in an instrument update between two icIEF instruments, ICE3 (original instrument, protein simple, bio-techne) and Maurice C (new instrument, protein simple, bio-techne). As study material, reference standard material of a globular glycoprotein and the respective icIEF analytical procedure were used.

Samples and matrix were prepared immediately before analysis. The matrix, containing urea, methyl cellulose, phosphoric acid as anode spacer, carrier ampholytes and two pI markers, was set up as a master mix and added to each sample of the respective sequence. Samples were detected by UV absorbance at 280 nm. The fluorescence detection mode of Maurice C was not used as this would be a significant discrepancy from the original analytical procedure validated on ICE3. ICE3 only allows for UV absorbance detection.

Samples were mixed with matrix, pipetted into 96-well plates for high-throughput analysis (046-021, protein-simple, bio-techne) and injected into Maurice iCIEF Cartridges (PS-MC02-C, protein simple, bio-techne), set up with catholyte and anolyte solutions by the manufacturer (#102506, protein simple, bio-techne).

Peak integration of the product was evaluated in three regions (region 1 – acidic region, region 2 – intermediate region, region 3 – basic region), each consisting of a sum of multiple peaks.

System suitability was assessed on Maurice C with the Maurice cIEF System Suitability Kit (#046-044, protein simple, bio-techne) following the manufacturer’s instructions and criteria. The peptide panel provided in the kit was analyzed in a bracketing approach at the beginning and end of each sequence. All experiments passed the system suitability test (SST).

Maurice C instruments were operated with the Empower™ 3 software (Waters™). For setup and calibration of Maurice C instruments, 0.5% methyl cellulose (#102730, protein simple, bio-techne) and Maurice CIEF Fluorescence Calibration Standard (#046-025, protein simple, bio-techne) were used according to the manufacturer’s recommendation.

Collected data were analyzed and integrated in Empower™ 3 software (Waters™). Results were evaluated in Microsoft Office Excel (Version 2308) and Minitab Statistical Software (Minitab® 20.2, 64-bit). For visual evaluation, electropherogram figures of both instruments were generated in Empower™ 3 (Waters™) and overlayed in Microsoft Office Excel.

Acceptance criteria were either visual (baseline comparability, resolution), or adopted from the original method validation on ICE3 (LOQ, response), or calculated from control chart data of historical measurements on ICE3 (peak positions in x, relative peak areas, measurement variance). The applied control chart data were collected from the same reference standard over four years, on three ICE3 instruments, by seven analysts, using multiple capillaries and generated a total of 128 data points. Acceptance criteria were identified / calculated as described in paragraph 2 Theory.

## 4 Results and discussion

### 4.1 Study design

The study design allows to generate and evaluate results for systematic instrument comparison. The results may either provide a detailed understanding of potential discrepancies between instruments, allowing to rationalize the planning of a partial or full method re-validation. Or, in case comparability can be inferred, the data may be a scientific foundation to move analytical procedures seamlessly to the new instrument. In this case, analytical procedures would be maintained and benefit from the advantages of more modern instruments, such as facilitated handling, higher standards in data integrity or more technical options, e.g. during trouble shootings.

### 4.2 icIEF instrument comparability study

Acceptance criteria for icIEF instrument comparability were generally an equal or better performance of Maurice C. The results are summarized in Figure 3 and Figure 4 and explained in more detail in the following paragraphs.

**Figure 3.**
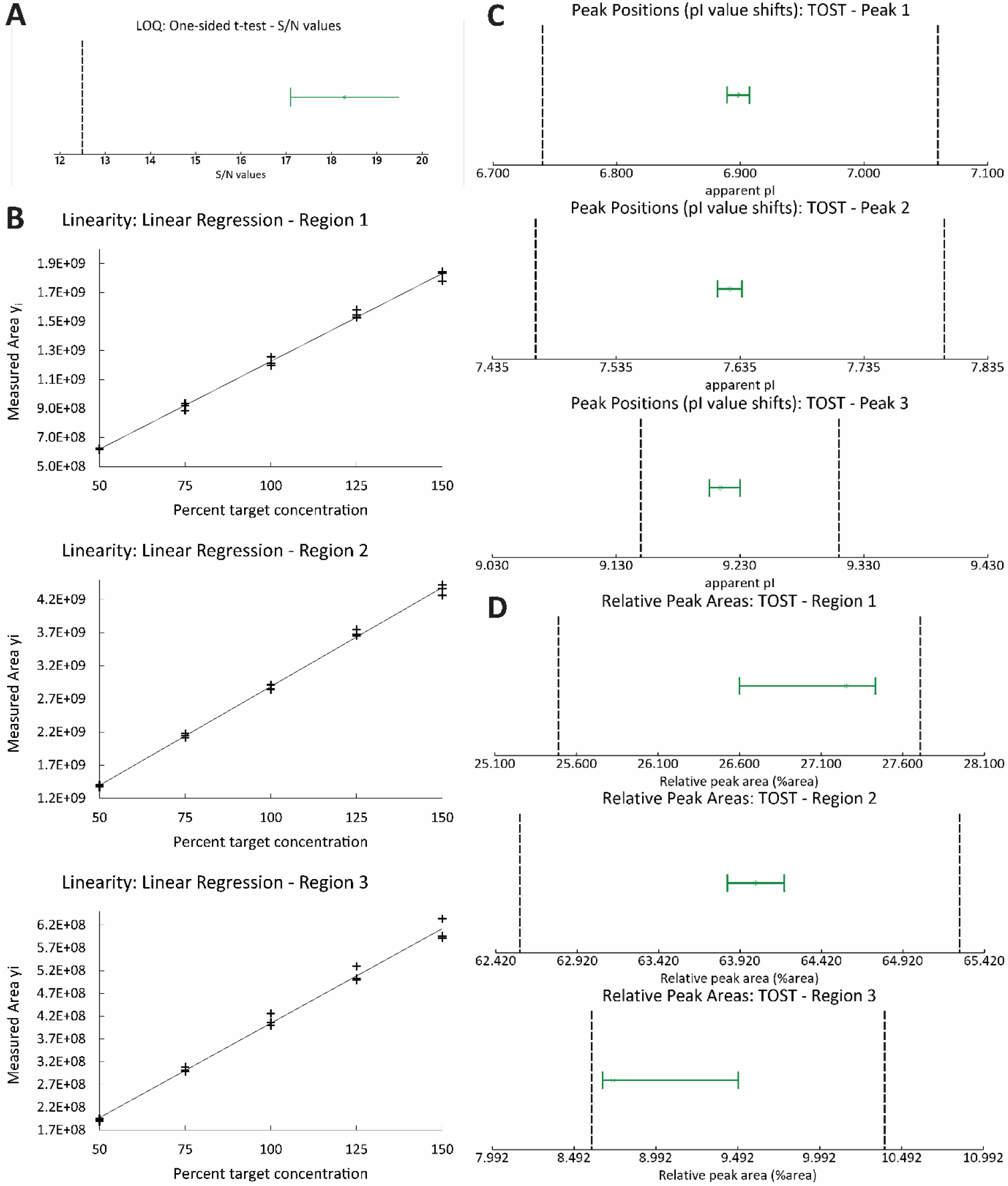
Statistical evaluation of icIEF instrument bridging study results. (A) LOQ: T-test of LOQ-peak S/N values with a 95% confidence interval (green interval bar) and the lower limit (black dashed line, S/N=12.5) received from original method validation on ICE3, (B) Linearity: Linear regression plots of measured peak areas plotted against percent target concentration. One subpanel per evaluated region (1, 2 and 3) (C) Peak Position shifts in x: TOST of apparent pI values with a 95% confidence interval for equivalence (green interval bar). Lower and upper equivalence limits (black dashed lines) were calculated from control chart data (mean ± 3SD range). (D) Relative Peak areas: TOST of relative peak areas with a 95% confidence interval for equivalence (green interval bar). Lower and upper equivalence limits (black dashed lines) were calculated from control chart data (mean ± 3SD range).

**Figure 4.**
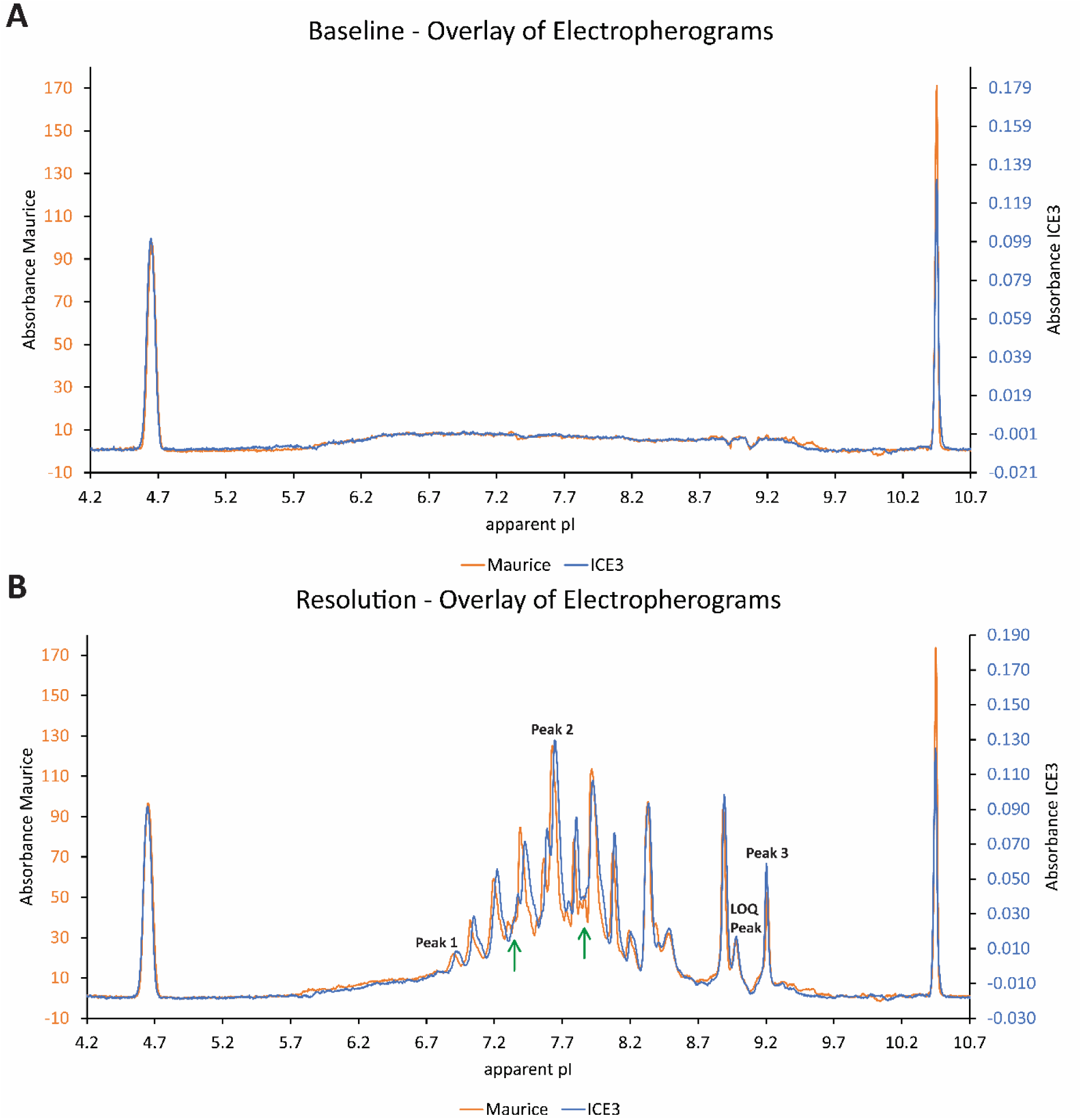
Visual evaluation of icIEF instrument bridging study results. X-Axes: Absorbance values in arbitrary units for Maurice (orange) and ICE3 (blue). Y-Axes: apparent pI values. (A) Baseline: Representative overlay of blank electropherograms (orange – Maurice C., blue – ICE3). (B) Resolution: Representative overlay of sample electropherograms (orange – Maurice C., blue – ICE3). Green arrows indicate discrepancies in resolution. Black labels indicate peaks used for evaluation of peak position shifts in x.

#### 4.2.1 Limit of signal quantitation

The limit of signal quantitation was according to the original method validation determined from a small peak in the electropherogram (Figure 4). Resolution discrepancies between the two instruments (higher resolution on Maurice C, see paragraph 4.2.5) caused the nominal LOQ-value determined on Maurice C to be higher (1.5%Area) than on ICE3 (1.2%Area). However, the S/N on Maurice C was non-inferior to ICE3 data (Mean(S/N Maurice C) = 18.3, Mean(S/N ICE3) = 12.7), as was confirmed with a one-sided t-test (95% confidence interval) (Figure 3A). Furthermore, the peak was more robust on Maurice C (RSD (%Area) = 6.2%) than on ICE3 (RSD (%Area) = 9%). Thereby, comparability of the Maurice C instrument with respect to the LOQ was confirmed.

#### 4.2.2 Response between product concentration and observed signal

The response was determined according to the procedure of the original method validation in a linearity experiment over a range of 50-150% target concentration with five equidistant concentration levels and three independent preparations (n=3) per level. Linear regression of product concentration and peak area was determined, and the correlation coefficient calculated for the total peak area (Figure S1) and the individual regions 1, 2 and 3 (Figure 3B). Acceptance criterion for comparability was according to the original method validation a correlation coefficient of r≥0.98. Values from original validation on ICE3 were 0.990 (Region 1), 0.990 (Region 2) and 0.990 (Region 3), newly measured values on Maurice C were 0.998 (Region 1), 0.999 (Region 2) and 0.996 (Region 3). Thereby, comparability of the Maurice C instrument with respect to response was confirmed.

#### 4.2.3 Baseline comparability

Baseline comparability was assessed visually by overlaying three blank electropherograms of Maurice C measurements with randomly selected historical ICE3 measurements (Figure 4A, Figure S2A and B). Visual evaluation showed equal occurrence of irregularities in the blank shape (e.g. increasing signal between apparent pI 5.7 and 6.5; indentations at apparent pI 8.9 and 9.1). Minor discrepancies in the position of irregularities in x could be observed (apparent pI 9.3 to 9.5) and were traced back to the usage of different carrier ampholyte batches, which can minimally affect the focusing behavior. No substantial discrepancies or deviating trends could be assessed in comparison of data from both instruments. Consequently, comparability of the Maurice C instrument with respect to baseline comparability was confirmed.

#### 4.2.4 Peak position shifts in x

Peak position shifts in x were assessed on three peaks located over a range of apparent pI 6.5 to 9.5 in the electropherogram (Figure 4B black labels). 12 data points (each being a mean value of six injections) from the matrix experiment were compared with historical control chart data collected on the ICE3 instrument (128 data points). Results are displayed in a blinded manner with respect to the exact pI values in Figure 3C. Equivalence of the data was tested using a TOST with a significance level of α=0.05 and could be confirmed for all peaks.

#### 4.2.5 Resolution

Resolution comparability was assessed visually by overlaying three exemplary electropherograms of Maurice C measurements with historical ICE3 measurements (Figure 4B, Figure S2C and D). Visual evaluation showed equal overall electropherogram appearance. Individual peak shapes displayed minimally higher resolution on Maurice C (see Figure 4B green arrows at apparent pI 7.2 to 7.4 or 7.8 to 8.0), which could either be traced back to the usage of different carrier ampholyte batches, which can minimally affect the focusing behavior or indicated a higher resolution (i.e. better focusing behavior) of Maurice C. Consequently, comparability of the Maurice C instrument with respect to resolution was confirmed.

#### 4.2.6 Relative peak area changes

Relative peak area changes were assessed on three regions of the electropherogram each a sum of multiple peaks. 12 data points (each being a mean value of six injections) from the matrix experiment were compared with historical control chart data collected on the ICE3 instrument (128 data points). Results are displayed in a blinded manner with respect to the exact relative peak area values in Figure 3D. Equivalence of the data was tested using a TOST with a significance level of α=0.05 and could be confirmed for all regions.

#### 4.2.7 Measurement variance

Measurement variance was assessed on three regions of the electropherogram each a sum of multiple peaks. Data from the matrix experiment comprising 12-fold sample preparation and measurement by two analysts on two instruments using four different cartridges on six days. Acceptance criteria were set by calculating the RSD% values of the relative peak areas of the three evaluated peak regions from historical control chart data collected on ICE3 (128 data points) and multiplying by a factor of 1.5 (same factor as used for intermediate precision acceptance criteria setting in original validation).

Criteria were RSD ≤ 2.1% (region 1), RSD ≤ 1.1% (region 2) and RSD ≤ 4.7% (region 3). Values from original validation on ICE3 were 0.7% (region 1), 0.4% (region 2) and3.2% (region 3), newly measured values on Maurice C were 1.5% (region 1), 0.6% (region 2) and 1.5% (region 3). RSD point values acquired from measurements on Maurice were below the acceptance criteria and thereby, comparability of the Maurice C instrument with respect to measurement variance was confirmed.

## 5 Concluding remarks

The presented study design is a comprehensive approach to rationalize the decision process of how to implement an instrument change in a GMP environment. It allows for a differentiated analysis of instrument comparability and thus provides a solid scientific fundament for maintenance of the analytical state-of-the-art in release analytics. It may well serve as a universal basis for instrument bridging approaches in various analytical methods.

The study design was successfully applied in the GMP environment of QC release analytics. The results demonstrate comparability of analytical results between two icIEF instruments and thus comprehensively complement a recent inter-company study initiated and published by the manufacturer protein simple [14] as well as available instrument comparability data by protein simple [15]. We additionally tested all other affected products on the new instrument in a lean matrix experiment (Figure 2, green box; data not shown), to evaluate transferability of product specific SST criteria. The SST criteria were met in all cases. The results of the comprehensive comparability study together with the passed SST criteria of all affected products present a solid scientific foundation for a seamless continuation of icIEF QC release analytics on the new instrument Maurice C.

## Supporting information

UniversalStudyDesign_Supplement_Ries

## Abbreviations

icIEF: imaged capillary isoelectric focusing
QC: Quality Control
GMP: Good Manufacturing Practice
TOST: two one-sided t-tests
DoE: Design of Experiments
SST: System suitability test

## Acknowledgements

The authors thank Gerald Knebl for valuable input on the study design, Dr Göran Hübner and Dr Heike Volkmer for valuable input to the manuscript and Dr Jakob Engel, Dr Beatrix Metzner, Dr Dominik Conrad, Dr Rainer Ilg, Volker Selzle and Gerald Knebl for input on the current status and regulations regarding life cycle instrument updates in the field of pharmaceutical release analytics.

The authors thank Anne Eberhardt, Dr Kerstin Kojer, Andreas Saumweber, Dr Ute Rockinger, Dr Jannik Brückmann, Jana Engler, Heidi Stark, Sabine Sautter, Alexander Wegner, and Milena Maier for their support during implementation of the icIEF instrument bridging in the Boehringer Ingelheim Biopharma QC environment.

## Author contributions

Conceptualization, F.H., F.T.W.; Methodology, A.B.R., F.H., M.N.M., F.T.W.; Validation A.B.R., M.N.M., F.T.W.; Formal analysis, A.B.R., D.K., M.N.M.; Investigation A.B.R., M.P., K.C., R.G., M.N.M.; Resources, F.H., S.E., F.T.W.; Data Curation, R.G., K.C., M.P., A.B.R., M.N.M.; Writing - Original Draft, A.B.R.; Writing – Review & Editing, F.T.W., M.N.M., D.K., S.E., A.B.R.; Visualization, A.B.R.; Supervision F.T.W.; Project administration, A.B.R.

## Data Availability Statement

The data that support the findings of this study are available from the corresponding author upon reasonable request.

## Supporting information

**<u>Supporting information file:</u>** UniversalStudyDesign_Supplement_Ries.docx

