## Supplementary material for "Universal Study Design for Instrument Changes in Pharmaceutical Release Analytics": UniversalStudyDesign_Supplement_Ries

### Supporting Information to "Universal Study Design for Instrument Updates in Pharmaceutical Release Analytics"

**Table S1** Matrix setup for measurement variance determination (Kojima design [13])

| Day | Analyst | Instrument | Capillary | No. of sample preparations |
|---|---|---|---|---|
| 1 | A | A | A | 2 |
| 2 | A | B | A | 2 |
| 3 | A | A | B | 2 |
| 4 | B | B | A | 2 |
| 5 | B | A | B | 2 |
| 6 | B | B | B | 2 |

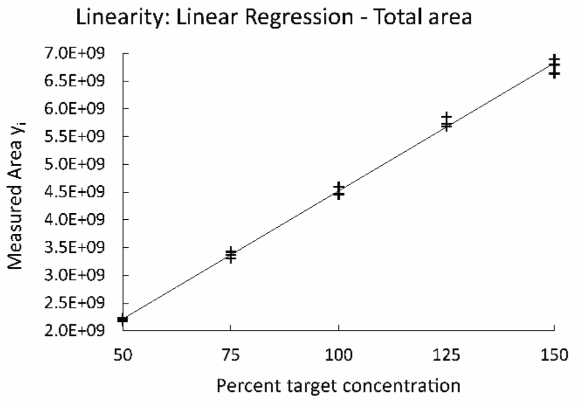


**Figure S1** Linearity: Linear regression plot of total peak area plotted against percent target concentration.

Linearity results: Correlation coefficient of total area from original validation on ICE3 is 0.99, newly measured value on Maurice is 0.999.

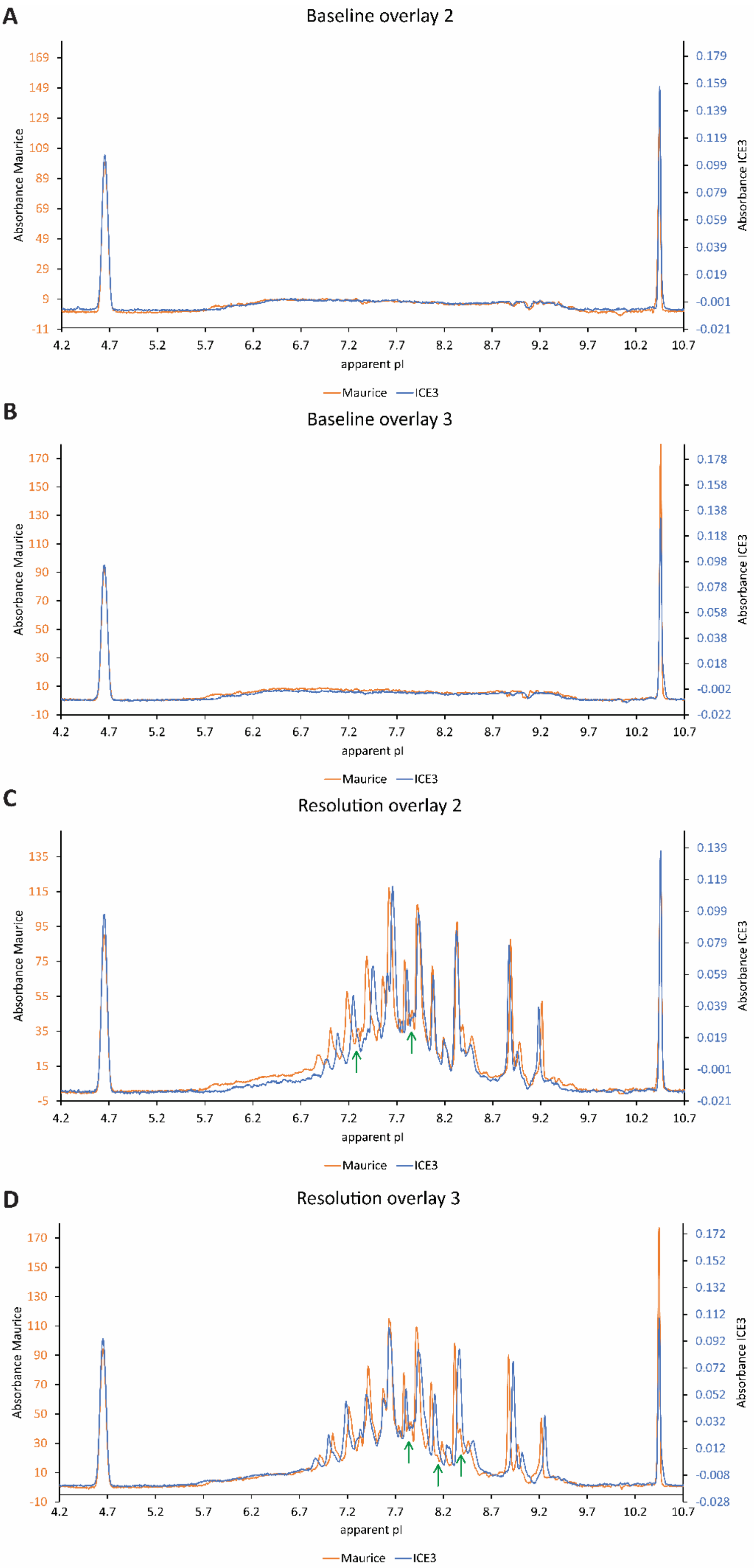
A
Baseline overlay 2
Absorbance Maurice
169
149
129
109
89
69
49
29
9
-11
Absorbance ICE3
0.179
0.159
0.139
0.119
0.099
0.079
0.059
0.039
0.019
-0.001
-0.021
4.2 4.7 5.2 5.7 6.2 6.7 7.2 7.7 8.2 8.7 9.2 9.7 10.2 10.7
apparent pI
Maurice ICE3
B
Baseline overlay 3
Absorbance Maurice
170
150
130
110
90
70
50
30
10
-10
Absorbance ICE3
0.178
0.158
0.138
0.118
0.098
0.078
0.058
0.038
0.018
-0.002
-0.022
4.2 4.7 5.2 5.7 6.2 6.7 7.2 7.7 8.2 8.7 9.2 9.7 10.2 10.7
apparent pI
Maurice ICE3
C
Resolution overlay 2
Absorbance Maurice
135
115
95
75
55
35
15
-5
Absorbance ICE3
0.139
0.119
0.099
0.079
0.059
0.039
0.019
-0.001
-0.021
4.2 4.7 5.2 5.7 6.2 6.7 7.2 7.7 8.2 8.7 9.2 9.7 10.2 10.7
apparent pI
Maurice ICE3
D
Resolution overlay 3
Absorbance Maurice
170
150
130
110
90
70
50
30
10
-10
Absorbance ICE3
0.172
0.152
0.132
0.112
0.092
0.072
0.052
0.032
0.012
-0.008
-0.028
4.2 4.7 5.2 5.7 6.2 6.7 7.2 7.7 8.2 8.7 9.2 9.7 10.2 10.7
apparent pI
Maurice ICE3

**Figure S2** Visual evaluation of icIEF instrument bridging study results. (A) Baseline: Second overlay (orange – Maurice C., blue – ICE3). (B) Baseline: Third overlay (orange – Maurice C., blue – ICE3). (C) Resolution: Second overlay (orange – Maurice C., blue – ICE3). Green arrows indicate discrepancies in resolution. (D) Resolution: Third overlay (orange – Maurice C., blue – ICE3). Green arrows indicate discrepancies in resolution.
